# To B or not to B: *Arsenophonus* as a source of B-vitamins in whiteflies

**DOI:** 10.1101/311076

**Authors:** Diego Santos-Garcia, Ksenia Juravel, Shiri Freilich, Einat Zchori-Fein, Amparo Latorre, Andrés Moya, Shai Morin, Francisco J. Silva

## Abstract

Insect lineages feeding on nutritionally restricted diets such as phloem, xylem, or blood, were able to diversify by acquiring bacterial species that complemented the missing nutrients. These bacteria, considered obligate/primary endosymbionts, share a long evolutionary history with their hosts. In some cases, however, these endosymbionts are not able to fulfill all the nutritional requirements of their host, driving the acquisition of additional symbiotic species. Whiteflies, which feed on phloem, established an obligate relationship with *Candidatus* Portiera aleyrodidarum, who provides essential amino acids and carotenoids to the host. As many Whiteflies species harbor additional endosymbionts, they could provide their hosts with missing nutrients. To test this hypothesis, genomes of several endosymbionts from the whiteflies *Aleurodicus dispersus*, *A. floccissimus* and *Trialeurodes vaporariorum* were sequenced and analyzed. All three species were found to harbor an endosymbiont from the genus *Arsenophonus*, and the two former also host *Wolbachia*. A comparative analysis of the three *Arsenophonus* genomes revealed that although all of them are capable of synthesizing B-vitamins and cofactors, such as pyridoxal, riboflavin, or folate, their genomes and phylogenetic relationship vary greatly. *Arsenophonus* of *A. floccissimus* and *T. vaporariorum* belong to the same clade and display characteristics of facultative endosymbionts, such as large genomes (3 Mb) with thousands of genes, many pseudogenes, intermediate GC content, and mobile genetic elements (MGEs). In contrast, *Arsenophonus* of *A. dispersus* belongs to a different lineage and displays the characteristics of a primary endosymbiont, such as a reduced genome (670 kb) with 400 genes, 32% GC content, and no MGEs. However, the presence of 274 pseudogenes suggests that this symbiotic association is more recent than other reported hemipteran’s primary endosymbionts. *Arsenophonus* of *A. dispersus* gene repertoire is completely integrated in the symbiotic consortia, and the biosynthesis of most vitamins occurs in shared pathways with its host. In addition, *Wolbachia* have also retained the ability to produce riboflavin, FAD, and folate, and may have a nutritional contribution. Taken together, our results show that *Arsenophonus* have a pivotal place in whiteflies nutrition by their ability to produce B-vitamins, even if they diverge and/or go through a genome reduction process.

## 2 Introduction

Diverse eukaryotes live together with bacterial symbionts. Mutualistic, commensal, or parasitic relationships have been described for these associations (Moya et al., 2008). The Class Insecta is an example of mutualistic associations and several lineages are known to live in intimate relationship with an obligate symbiont for millions of years. Such symbionts are usually harbored inside specialized cells, termed bacteriocytes, and are vertically transmitted from the mother to her offspring. These bacterial symbionts, living inside cells and transmitted for long evolutionary periods, has been denominated primary (or obligatory) endosymbionts. The evolution of these symbioses had several effects on the gene repertoires of both hosts and endosymbionts, but is most notedly characterized by drastic reductions in the latter (Latorre and Manzano-Marín, 2016). Host diet complementation has frequently initiated the symbiosis, but in some cases, the host has nutritional requirements which the primary endosymbiont is unable to satisfy, and secondary (or facultative) endosymbiont are thus required for diet complementation. When these requirements became vital, it is possible that a secondary endosymbiont becomes primary, resulting in a consortium of primary endosymbionts.

Whiteflies and psyllids are two sister hemipteran lineages, which probably started their evolutionary success through the ancient association with ancestral bacterial species of the family Halomonadaceae (Santos-Garcia et al., 2014a). This bacterial lineage splits into *Candidatus* Portiera aleyrodidarum (hereafter, the term Candidatus will be used only at the first time a species is mentioned) in whiteflies and *Candidatus* Carsonella ruddii in psyllids. Based on gene repertoires, it was proposed that *Portiera* collaborates with its hosts in the synthesis of amino acids and carotenoids (Santos-Garcia et al., 2012, 2015; Sloan and Moran, 2012). Because whiteflies feed only on the phloem sieve-element, their diet is also deficient in vitamins and cofactors. Individual whiteflies may carry one or several additional endosymbionts belonging to seven genera (Zchori-Fein et al., 2014). It has been proposed that some of these additional endosymbionts are likely involved in vitamins and cofactors supply to their host, as for example, *Candidatus* Hamiltonella defensa (Rao et al., 2015). *Candidatus* Arsenophonus is a genus of secondary endosymbiont that has been observed in many whitefly species (Thao and Baumann, 2004; Zchori-Fein et al., 2014; Santos-Garcia et al., 2015). Similar to *Hamiltonella*, it is only found inside bacteriocytes, while other endosymbionts like *Candidatus* Wolbachia sp. or *Candiddatus* Rickettsia sp. present different tissue tropisms (Gottlieb et al., 2008; Skaljac et al., 2010, 2013; Marubayashi et al., 2014). *Arsenophonus* lineages have also evolved intimate associations with other insect taxa with different diets, such as blood-sucking insects, where the supplementation of B-vitamins and cofactors were their proposed role (Šochová et al., 2017).

Whiteflies (Sternorrhyncha:Aleyrodidae) are composed mainly by two subfamilies, the Aleurodicinae and the Aleyrodinae. These two subfamilies probably originated between the Jurassic (Drohojowska and Szwedo, 2015; Shcherbakov, 2000) and the Middle Cretaceous (Campbell et al., 1994). While the Aleyrodinae contains the largest number of species described so far (140 genera, 1440 species), the Aleurodicinae has a relatively low number of described species (17 genera, 120 species) (Ouvrard and Martin, 2018). Analysis of the genome content of *Portiera* from the whitefly species *Aleurodicus dispersus* and *A. floccissimus* (Aleurodicinae) and *Trialeurodes vaporariorum* (Aleyrodinae) suggested that they are unable to complement their host diet with essential cofactors and vitamins (Santos-Garcia et al., 2015). The work presented here was thus initiated in order to test the prediction that *Arsenophonus* and/or *Wolbachia*, the other symbionts present in these species, are capable of providing their hosts with the lacking cofactors and vitamins.

## 3 Material and Methods

### Sequences retrieval

Sequences of bacteria other than *Portiera* were retrieved from a previous shotgun sequencing project of three whiteflies species: *A. dispersus*, *A. floccissimus*, and *T. vaporariorum* (Santos-Garcia et al., 2015). In brief, *T. vaporariorum* was collected in 2014 near the IRTA Institute of Agrifood Research and Technology (Barcelona, Spain) and identified by Dr. Francisco JosÃ© Beitia. *A. dispersus* and *A. floccissimus* were collected and identified by Dr. Estrella Hernandez Suarez in 2014 from banana fields (Tenerife Island, Spain). All three whiteflies harbored *Portiera*, *Arsenophonus* and *Wolbachia* endosymbionts. Genomic DNA (gDNA) was obtained using and alkaline lysis method from single bacteriomes (dissected with glass micro-needles). For each species, ten single bacteriome gDNA extractions were subjected to a Whole Genome Amplification process (WGA; GenomiPhi V2, GE Healthcare) and pooled by species. WGA gDNAs were sequenced by Illumina HiSeq 2000 using a mate-paired library (100bp*2, 3kb insert size). A full description can be found at Santos-Garcia et al. (2015).

### Metagenome-assembled genomes (MAGs)

For each species library, RAW Illumina reads were quality checked, and trimmed/clipped if necessary, with FastQC v0.11.3 (Andrews, 2010) and TrimmomaticPE v0.33 (Bolger et al., 2014). Kraken v0.10.6 (Wood and Salzberg, 2014) was used to classify the RAW Illumina reads using a custom genomic database including: *Portiera*, *Hamiltonella*, *Rickettsia*, *Wolbachia*, *Arsenophonus*, *Candidatus* Cardinium hertigii, several whiteflies’ mitochondria, *Bemisia tabaci* MEAM1, and *Acyrthosiphon pisum* (Table S1). Reads classified as mitochondrial, insect or *Portiera* were discarded. The rest of reads were assembled with SPADES v3.11.0 (-meta -careful -mp) (Nurk et al., 2017). Obtained contigs were classified using Kraken and the custom database. Contigs belonging to known whiteflies endosymbionts were recovered and added to the custom database. Trimmed/clipped reads were re-classified and reads belonging to known whitefly endosymbionts were re-assembled alone with SPADES (-careful -mp) to produce metagenome-assembled genomes (MAGs). Illumina reads were digitally normalized (khmer v1.1) before the re-assembly stage (Crusoe et al., 2015). MAGs contigs were scaffolded and gap-filled with SSPACE v3 (Boetzer et al., 2011) and Gapfiller v1.10 (Boetzer and Pirovano, 2012), respectively. Finally, a manual iterative mapping approach, using Illumina trimmed/clipped reads, was performed with Bowtie2 v2.2.6 (Langmead and Salzberg, 2012), MIRA v4.9.5 (mapping mode) (Chevreux, B., Wetter, T. and Suhai, 1999), and Gap4 (Staden et al., 2000) until no more contigs/reads were joined/recovered for each MAG. Finally, Bowtie2 and Pilon v.1.21 (-jumps) (Walker et al., 2014) with Illumina reads were used to correct MAGs contigs.

### MAGs annotation

MAGs initial annotations were performed with prokka v1.12 (Seemann, 2014). Enzyme commission numbers were added with PRIAM March 2015 release (Claudel-Renard et al., 2003). Gene Ontology, PFAM, and InterPro terms were added with InterProScan v5.27-66 (Jones et al., 2014). Putative pseudogenes and their positions in the genome were detected with LAST using several bacterium relative proteomes as a query (Kielbasa et al., 2011). Insertion Sequences (IS) were predicted with ISsaga (Varani et al., 2011). *B. tabaci* genome annotations (GCA 001854935.1) were downloaded and refined with PRIAM and InterProScan. MAGs, their corresponding *Portiera* (Santos-Garcia et al., 2015) and *B. tabaci* metabolism comparisons were performed on PathwayTools v21.5 (Karp et al., 2002). Heatmap and clustering was performed on R (R Core Team, 2018) with ggplots2 (Wickham, 2009).

### Comparative genomics

OrthoMCL v2.0.9 was used to compute clusters of orthologous proteins (Li et al., 2003). Cluster of Orthologous Groups (COGs) terms were assigned using diamond v0.8.11.73 (July 2016 bacterial RefSeq database, Buchfink et al. (2015)) and MEGAN6 (Huson et al., 2016). Average Nucleotide Identity (ANI) and Average Amino Acid Identity (AAI) values were obtained with the Enveomics tools (Rodriguez-R and Konstantinidis, 2016). Synteny between MAGs was checked with Mummer v3 (Kurtz et al., 2004) and Mauve, using *Arsenophonus* ARAD as reference for contig re-ordering (Darling et al., 2010). GenoPlotR was used to plot Mauve results (Guy et al., 2010).

The presence of potential orthologs of *B. tabaci* genes in the other whiteflies species was assessed using the Illumina cleaned data and diamond, with a blastx search strategy (minimum alignment length of 25 bp and 1*e^−3^* e-value) against the selected *B. tabaci* proteins (Table S2). Recovered reads were classified with Kraken, using the custom and the mini-kraken database, to discard bacterial reads. Finally, non-bacterial reads were used as query for a blastx search against the nr database (last access: March 2018) (Altschul et al., 1990) and their best hit classified with MEGAN6. Only reads assigned to the phylum “Arthropoda” were considered as genomic reads of the screened genes in the other whiteflies species.

### Phylogenetics

General phylogenetic position of the newly sequenced *Arsenophonus* and *Wolbachia* was assessed using several 16S rRNA genes downloaded from the GenBank. Genes were aligned with ssu-align (default masking) (Nawrocki, 2009). IQ-TREE was used to select the best substitution model (TVM+F+R3 and TN+F+R2, respectively) and to infer the majority rule maximum likelihood (ML) tree, with its associated support values (Nguyen et al., 2015; Kalyaanamoorthy et al., 2017).

Phylogenomic trees were generated using 82 and 97 conserved proteins, selected using PhyloPhlAn v0.99 (Segata et al., 2013), from *Arsenophonus* and *Wolbachia* inferred proteomes, respectively. Protein files were sorted by species name, using fastasort from exonerate v2.2.0 (Slater and Birney, 2005) and aligned with MAFFT v7.215 (Katoh et al., 2002). Proteins alignments were concatenated by species index using Geneious version 11.1.2 (Kearse et al., 2012). IQ-TREE was used to infer majority rule ML tree, and associated support values, under the substitution models JTTDCMut+F+R3 for *Arsenophonus* and JTT+F+I+G4 for *Wolbachia*.

## 4 Results

### Genomic features and comparative genomics

Five MAGs, two *Arsenophonus* and two *Wolbachia* from *A. dispersus* (ARAD, WBAD) and *A. floccissimus* (ARAF, WBAF) and one *Arsenophonus* from *T. vaporariorum* (ARTV) were recovered (Table 1). All MAGs were in a draft status but with different quality. Only ARAD, was assembled as a circular scaffold, supported by some mate-paired reads. However, a gap at the contig edges was not closed due to the presence of a *dnaX* duplication and a long AT low complexity region. The assembly of ARAF genome was also of high quality, with only 11 contigs and a N50 value higher than 500 kb. The rest of recovered MAGs were in different draft status, being the *Wolbachia* the most fragmented ones. Although *Wolbachia* was previously detected by PCR in *T. vaporariorum* (Santos-Garcia et al., 2015), its genome was impossible to assemble and analyze due to the low amount of reads recovered (Table S3). Indeed, only 21 *Wolbachia* contigs were recovered. From them, only one contig was above 1 kb (2.5X coverage) while the rest were below 400 nt. This suggests that *Wolbachia* is present in *T. vaporariorum* but at low amounts.

**Table 1:**
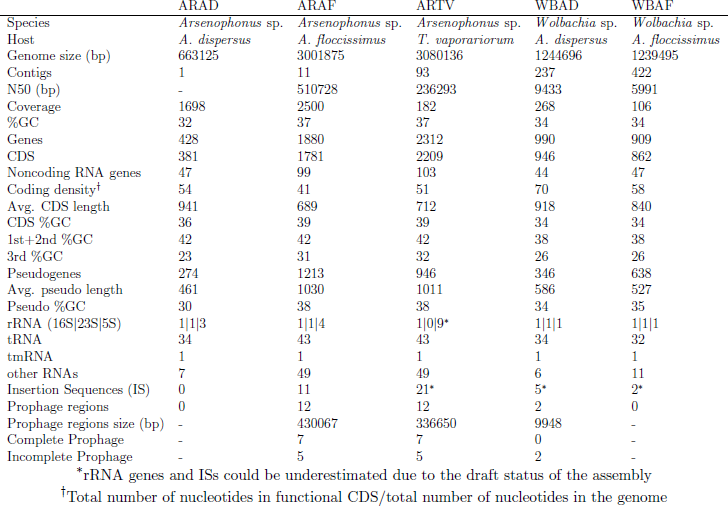
Assembly statistics and genomic features of the five endosymbionts sequenced.

According to their genomic features, the *Arsenophonus* sequenced can be divided into two groups. While ARAD displays typical characteristics of primary endosymbionts, ARAF and ARTV display genomic features closer to facultative or secondary endosymbionts (Table 1). This pattern is emphasized by the different genome sizes (0.67 *vs.* 3 Mb), their GC content (32 *vs.* 37%), the number of genes (428 *vs.* around 2000), and the presence of insertion sequences (ISs) and prophages (0 *vs.* 12). However, they have in common the low coding density (range 41 to 54%) due to the presence of almost as many pseudogenes as genes. Most of the prophages in ARAF and ARTV were unrelated (based on their ANI values) except the pairs of prophages 2/8 and 3/7 from ARAF and the pair of prophage 9 from ARAF and prophage 7 from ARTV (Figure S1, Figure S2 and Table S4).

Regarding *Wolbachia*, both recovered MAGs display sizes and genomic features similar to other sequenced species from this genus (Ellegaard et al., 2013), with WBAF being more fragmented and lacking two tRNAs. Two incomplete prophages were detected in the genome of WBAD (Figure S1).

### Phylogenetic placement

The phylogenetic position of each assembled *Arsenophonus* within the genus was tested using the 16S rRNA phylogeny, phylogenomics and ANI/AAI analyses. Based on 16S rRNA phylogenetic analysis, ARAD belongs to a different lineage than ARAF and ARTV (Figure S3). While ARAD was placed in a not well supported cluster, including *A. nasoniae* and *Arsenophonus* from other insects (the aphid *Stomaphis* and the dipteran *Ornithomya*), ARAF and ARTV clustered with endosymbionts of several whitefly species and other insects. The use of a species threshold of 95% ANI (Konstantinidis and Tiedje, 2005) suggests that *A. triatominarum*, *A. nilaparvata*, *A.* of *Entylia carinata*, ARAF and ARTV belong to the same species, whereas *A. nasoniae*, *A. lipopteni* and ARAD form three different species (Table S4, Figure S4). The AAI analysis showed the presence of two clusters, one for ARAD and *A. nasoniae* and the other for the remaining species, except the endosymbiont *A. liptoteni*, with a value smaller than 80% AAI. Phylogenomics results show a partial congruence with ANI/AAI, placing *A. triatominarum* as the closest relative of ARAD (Figure 1A). The two endosymbionts form a sister monophyletic clade to all the others *Arsenophonus*, which form a second monophyletic clade. *A. nasoniae* was placed at the basal position of both clades.

**Figure 1:**
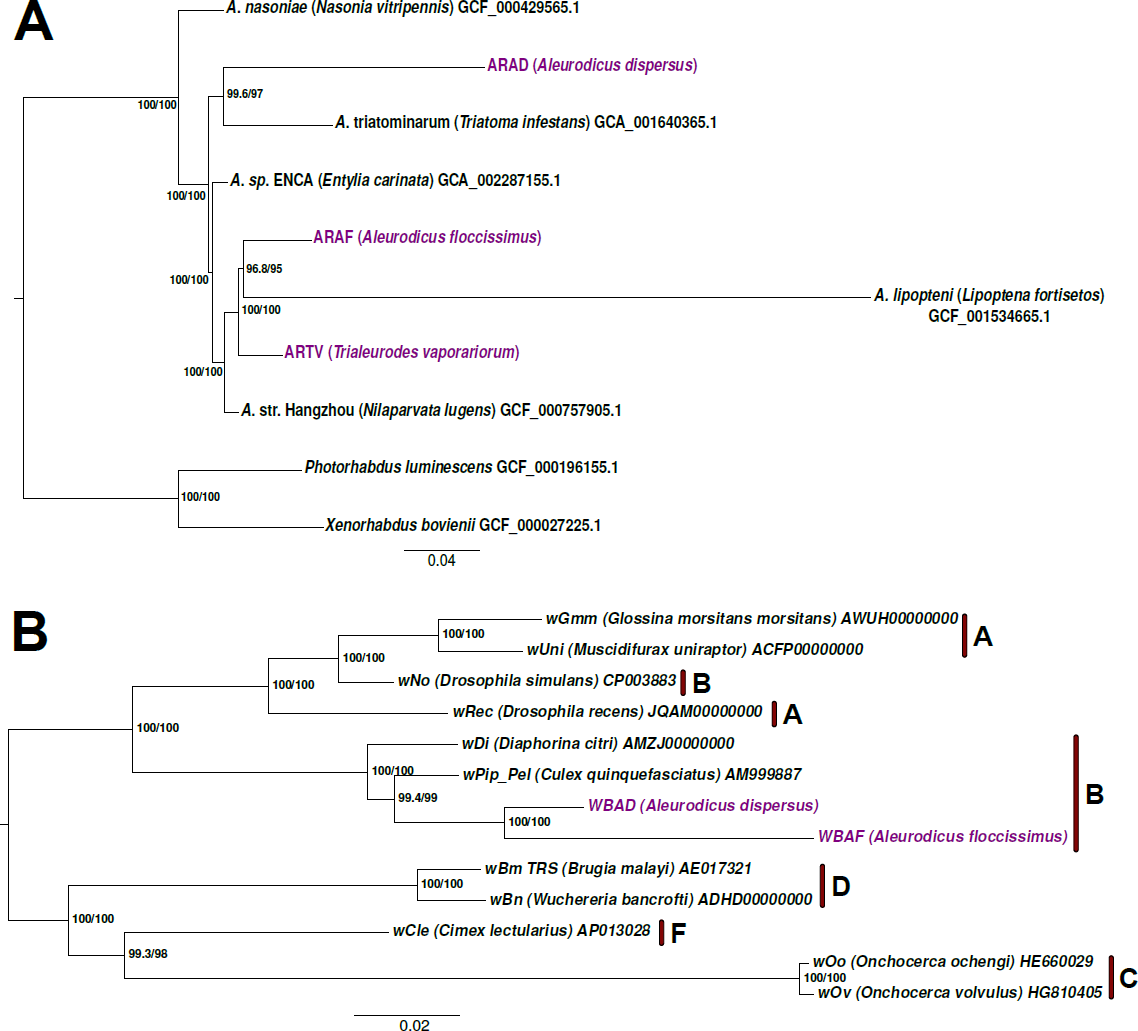
Phylogenomic trees of Arsenophonus and Wolbachia. (A) *Arsenophonus* (rooted) and (B) *Wolbachia* (unrooted) ML trees were inferred using 82 and 97 concatenated conserved protein alignments under a JTTDCMut+F+R3 and JTT+F+I+G4 substitution model respectively. Support values were obtained with 1000 ultrafast bootstraps (right node labels) and 5000 SH-aLRT (left node labels). Names for the eukaryotic hosts are shown after the strain names inside parentheses. Accession numbers are displayed for each of the proteomes used. *Arsenophonus* ARAD, ARAF, and ARTV and *Wolbachia* WBAD and WBAF are highlighted in purple. (B) Red bars and letters denote the different *Wolbachia* supergroups.

The phylogenetic positions of the two *Wolbachia* were tested with the same analyses. In the 16S rRNA phylogeny, WBAD (*A. dispersus*) and WBAF (*A. floccissimus*) form a well-supported clade inside a major clade, which contains *Wolbachia* strains from other whiteflies and other insect taxa (Figure S5). In addition, based on phylogenomics, WBAD and WBAF can be placed inside the *Wolbachia* B super group, with *Wolbachia* from *Culex quinquefasciatus* as the closest genome included in the analysis (Figure 1B). Finally, the high level of nucleotide identity (97.6% ANI value) suggests that both *Wolbachia*, WBAD and WBAF, are strains from the same species.

### Comparative genomics: synteny and functional categories

When the inferred proteomes from *Arsenophonus* ARAD, ARAF and ARTV were compared (Table S5), a core genome composed of only 289 clusters was obtained (Figure 2B). Interestingly, the core genome includes many clusters related to vitamins and cofactors biosynthetic pathways. The small number of shared protein clusters results from the reduced proteome of ARAD, which is mainly a subset of the larger ARTV and ARAF proteomes. Only 4 out of the 10 ARAD specific clusters were not hypothetical proteins: the pyruvate kinase II (pyruvate kinase I is present in the three proteomes), the 3-oxoacyl-[acyl-carrier-protein] synthase (*fabF*), which is involved in fatty acid and biotin biosynthesis, and proteins encoded by the genes *recA* and *nudE*. In case only ARAF and ARTV were compared, the core proteome would have contained more than 1,000 clusters. The number of species specific clusters of ARTV (997) and ARAF (609) were higher compared to ARAD, which is in accordance with their genome sizes. Synteny analysis highlights the strong genome reduction underwent in ARAD compared to ARAF and ARTV (Figure 2A). This includes the loosing of macrosynteny (general genome architecture), while maintaining microsynteny (e.g. operons). In contrast, ARTV and ARAD still show high level of macrosynteny, although some rearrangements were observed (Figure 2C).

**Figure 2:**
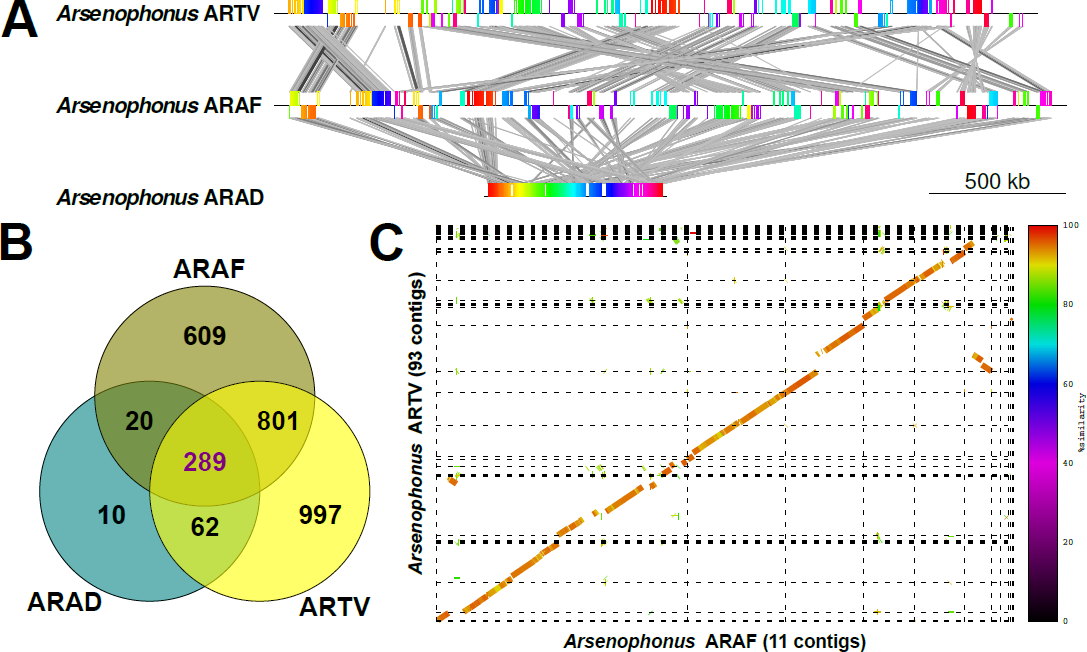
Comparative analyses of the three *Arsenophonus* MAGs. (A) Graphic linear representation of the genome rearrangements observed comparing ARTV with ARAF and ARAF with ARAD. Synteny blocks display the same color. ARAF and ARTV contigs were ordered according to ARAD before synteny blocks calculations with Mauve for a better visualization. (B) Venn diagram displaying the number of clusters found on each subspace of the *Arsenophonus* pangenome (Table S5). (C) Synteny between the draft genomes of ARTV and ARAF. Contig edges are displayed as dashed lines. Contigs of ARTV were ordered with Mauve using ARAF as reference for a better visualization.

*Wolbachia* ARAD and ARAF showed a core genome of 686 clusters, and 253 and 169 species specific clusters, respectively (Figure 3A, Table S6). It should be noted, however, that the highly fragmented status of these genomes could produce a large number of artificial specific clusters. Comparison of the two *Wolbachia* genomes revealed that some of the largest contigs show the same gene order and a high level of nucleotide identity (Figure 3B), suggesting that both macrosynteny and microsynteny are probably kept.

**Figure 3:**
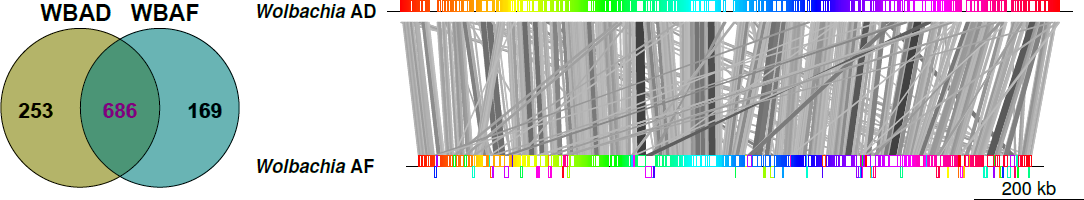
Comparative analyses of the two *Wolbachia* MAGs. (A) Venn diagram showing the shared and specific coding gene clusters between WBAD and WBAF (Table S6). (B) Synteny between the draft genomes (see Table 1) WBAD and WBAF. Contigs of WBAF were ordered with Mauve using WBAD as reference for a better visualization.

The proteomes of the five endosymbionts were functionally classified according to the Cluster of Orthologous Groups (COGs). Because of the strong reduction of the gene repertoire in *Arsenophonus* ARAD, this endosymbiont shows a smaller number of hits in each functional category, except J (translation and ribosomal structure and biogenesis), where the number of hits was similar to the other endosymbionts (Figure 4 left). In fact, in ARAD, this category has a relative frequency in the proteome higher than 0.20 4 right). In addition, category J and the other categories related to Information storage and processing (A, K, and L) showed higher relative frequencies in ARAD than in the other two *Arsenophonus* (Figure 4 right). Based on the number of hits and relative frequencies, the loss of the gene repertoire in categories G (carbohydrate metabolism), I (lipid metabolism), and O (post-translational modification and chaperones) was lower in ARAD. As expected, *Wolbachia* proteomes show the retention of the informational categories but also retain COGs related, among others, to environmental response.

**Figure 4:**
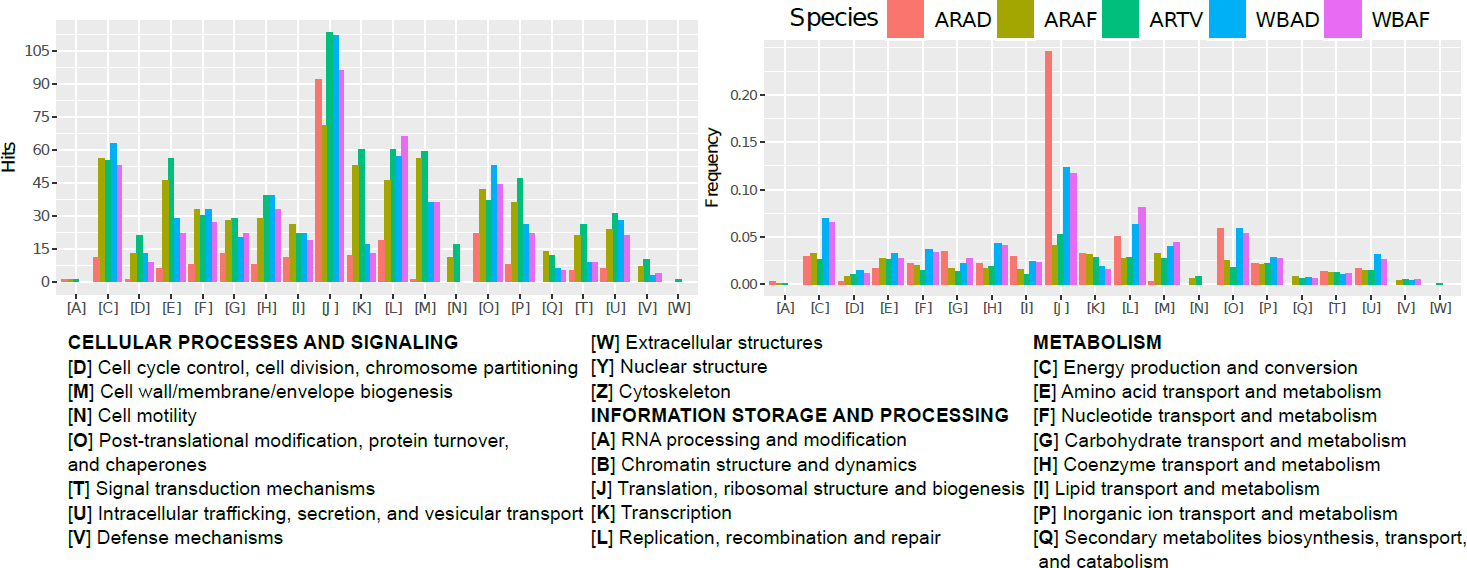
Distribution of the endosymbiont proteomes in functional categories. Bar plots showing the number of hits (left) and its relative frequency (right) in each functional category (COG) in the proteomes of the three analyzed *Arsenophonus* and the two analyzed *Wolbachia* MAGs.

### Biosynthetic metabolic potential

An integrated analysis of the metabolism of the whitefly bacteriocytes of the three species was performed in order to predict the potential biosynthetic capabilities of the harbored *Arsenophonus*, *Wolbachia* and *Portiera* (Santos-Garcia et al., 2015). That of the insect host was inferred using the *B. tabaci* genome (GCA 001854935.1). Amino acid biosynthesis in the three whiteflies is mainly conducted by the *Portiera* strains (PAAF, PAAD, and PATV), which maintain the ability to produce four (PATV)/five (PAAF and PAAD) out of ten essential amino acids, and require some complementation from the host for the synthesis of the others six (Figure 5A, cluster A). While some cofactors/vitamins and precursors are produced by almost all the endosymbionts and the host (cluster B), others are mainly produced by some *Arsenophonus* (cluster C), both *Arsenophonus* ARAF and ARTV, *Wolbachia* and the host (cluster D), or, apparently, by none of them (cluster E). In general, both *Wolbachia* present a biosynthetic potential that is lower than that of the most reduced *Arsenophonus*, ARAD. In addition, while both *Wolbachia* still present all the electron transport chain, *Arsenophonus* ARAF and ARTV have lost the Ubiquinol oxidase and ARAD only maintains the Cytochrome-c oxidase and has lost the ATP synthase (not shown).

**Figure 5:**
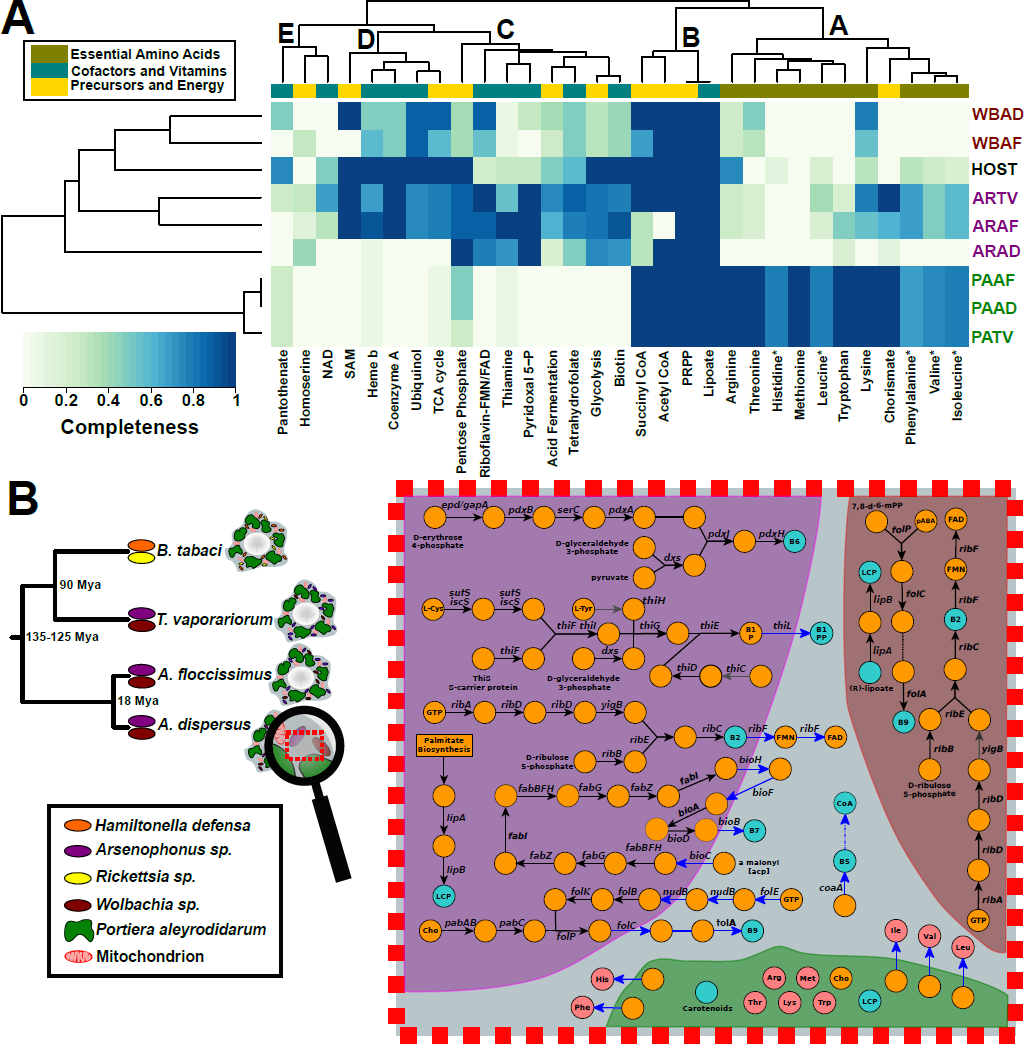
Biosynthetic potential of *Arsenophonus*, *Wolbachia*, *Portiera* and white-flies. (A) Heat map and hierarchical clustering showing, in a color scale, the degrees of completeness of several compounds’ biosynthetic pathways, including essential amino acids, cofactors and vitamins. Completeness was measured as number of enzymes present in the analyzed genome over the total number of enzymes in the pathway. *Arsenophonus* (AR), *Wolbachia* (WB) and *Portiera* (PA) from *A. dispersus* (AD), *A. floccissimus* (AF) and *T. vaporariorum* (TV) are highlighted in purple, red and green respectively. *B. tabaci* genome was used as a general host representative. (B) On the left, phylogenetic relationship and endosymbionts content of the *Aleurodicus*, *T. vaporariorum* and *B. tabaci* whiteflies (Santos-Garcia et al., 2015). Sequences from *Wolbachia* were scarcely detected in *T. vaporariorum*, suggesting low amounts of this bacterium in the insect. On the right, schematic representation of some vitamins and precursors metabolic pathways occurring at the bacteriocyte of *A. dispersus*. *Arsenophonus* ARAD is represented in purple, *Wolbachia* WBAD in redwood, *Portiera* in green and the host’s cytosol in blue. Black arrows denote enzymes present in the corresponding endosymbiont, blue arrows enzymes present in the host, grey arrows denote pseudo/absent genes, dashed arrows denote simplified pathways and dashed lines spontaneous reactions. Abbreviations: lipoyl carrier protein (LCP), chorismate (Cho), acyl carrier protein (acp), 4-aminobenzoate (pABA), (7,8-dihydropterin-6-yl)methyl diphosphate (7,8-d-6-mPP), diphosphate (PP). 13

Although the essential amino acid biosynthetic potential of *Arsenophonus* and *Wolbachia* is limited (Figure 5A), ARAF and ARTV still retain some genes involved in the synthesis of amino acids or their precursors, such as lysine (its precursor, meso-diaminopimelate is also involved in the peptidoglycan biosynthesis) and chorismate (the precursor of phenylalanine). In ARTV, the complete pathway, encoded by seven genes, from D-erythrose 4-phosphate and D-ribulose-5-phosphate to chorismate is functional (*aroF*, *aroB*, *aroQ*, *aroE*, *aroL*, *aroA*, and *aroC*). This, in combination with the presence of *tyrA*, *tyrB*, and an additional monofunctional chorismate mutase gene, suggests the ability of ARTV to produce tyrosine and a potential collaboration with *Portiera*, or the host, for the production of phenylalanine. The ability to synthetize amino acid was completely lost in ARAD, suggesting this not to be the reason why the lineage of ARAD evolved a close association with *A. dispersus*.

The contribution of the different partners to the synthesis of vitamins and cofactors is heterogeneous. ARTV and ARAF, the two *Arsenophonus* with larger genomes, are those maintaining higher capabilities, with the loss of several genes required for some of the pathways. Despite the reductive evolutionary process in ARAD that has drastically reduced its potentialities, some capabilities are still retained, suggesting that they might be important in the symbiotic relation (Figure 5A and 5B). ARAD, but also ARAF and ARTV, encode the complete pathway to synthesize pyridoxal 5’-phosphate (vitamin B6) and riboflavin (vitamin B2). However, while ARAF and ARTV are able to produce FMN and FAD because they harbor a functional *ribF* gene, it is pseudogenized in ARAD. Since this gene is present in all whitefly hosts, both flavin cofactors may be produced in *A. dispersus* through a complementation between ARAD and its host (Figure 5B, Table S2). Also, being complemented again by the host, ARAD could synthesize thiamine (vitamin B1) and folate (vitamin B9, from imported chorismate) (Figure 5B). Although ARAD has lost *thiH* gene, some host encoded proteins can replace its function (Table S2). ARAF and ARTV also require complementation by their host in order to produce folate. ARTV lost most of the pathway to produce thiamine, and the missing steps are not complemented by its host. Additionally, *B. tabaci* seems to be able to produce biotin using some horizontally transferred genes (*bioA*, *bioB*, and *bioD*) (Luan et al., 2015), a protein capable of replacing the BioC function, and the insect’s own fatty acid biosynthesis pathway. Genes with similar function to *bioABCDFH* genes were found to be present in *T. vaporariorum* genomic reads, suggesting that the whitefly is also capable of producing biotin by using only host genes. Because only genes similar to *bioBCFH* were identified in the genomic reads from both *Aleurodicus* species (Table S2), the presence of *bioA* and *bioD* in ARAD and ARAF is required for the cooperative synthesis of biotin. However, ARAD, unlike ARAF, has only retained the two missing genes in the genomes of both *Aleurodicus*.

In ARAF, the gene *yigB* which is part of the riboflavin pathway, is pseudogenized. This gene encodes for a Haloacid Dehalogenase (HAD) phosphatase, that belongs to a superfamily of enzymes. Therefore, the activity of the protein encoded by the gene, can be substituted by related genes in the endosymbiont or the host genomes (Manzano-Marín et al., 2015). Finally, all *Arsenophonus* conserve the glycolysis and the pentose phosphate pathways to produce the precursors required for riboflavin, thiamine, and pyridoxal. Indeed, in ARAD, the enzyme encoded by the *gatA* gene from the glycolysis pathway seems to have replaced the *epd* product in the pyridoxal pathway. The combination of *ilvC* encoded enzyme in *Portiera* (Price and Wilson, 2014) and the *panBC* genes that were horizontally transferred to *B. tabaci*, allows this whitefly to produce its own pantothenate (vitamin B5) and Coenzyme A. However, *panBC* genes are likely not present in the genomes of *A dispersus*, *A. floccissimus*, and *T. vaporariorum*, where only bacterial hits were recovered showing no similarity to the ones present in the *B. tabaci* genome. This suggests that these whiteflies are not able to produce pantothenate and they acquire it from the diet or other sources (Figure 5B and Table S2). However, failure on horizontal transferred genes detection could be an artifact from the methods used, since they are unable to discover new horizontally transferred genes in the whiteflies screened.

The third endosymbiont of *A. dispersus* and *A. floccissimus*, *Wolbachia* WBAD and WBAF, may be also contributing to the vitamin/cofactor synthesis of the system, as they have the complete pathway for riboflavin-FMN-FAD synthesis and potentially are able to produce folate from some intermediate precursors (Figure 5B).

## 5 Discussion

Some bacterial lineages show, predominantly, facultative symbiotic lifestyles associated with insects. In such cases, they can inhabit the cytoplasm of bacteriocytes and coexist with the primary endosymbiont. These bacterial lineages usually have large gene repertoires and genomes, allowing them to be autonomous and capable of infecting individuals from very different taxa. Examples of such clades are the genera *Arsenophonus*, *Sodalis*, and *Serratia* (Nováková et al., 2009; Manzano-Marín et al., 2017; Santos-Garcia et al., 2017). Because their presence may bare fitness costs to the host, they do not become fixed in the insect lineage except if their nutritional contribution (or other kind of benefit) is important and is required for periods of many generations. Under such a scenario, a facultative endosymbiont may evolve towards a more intimate association, becoming a co-primary symbiont (Lamelas et al., 2011). Although there are several features associated with this new lifestyle, the most relevant is the decrease in the genome size. In the lineage *Arsenophonus*-*Riesia*, the sizes of several genomes have been reported, ranging from the 0.58 Mb of *Candidatus* Riesia pediculicola (Kirkness et al., 2010) and 0.84 Mb of *A. lipopteni* (Nováková et al., 2016) to 3.86 Mb of *A. triatominarum*, but with several genomes of intermediate sizes (Šochová et al., 2017). Similarly, *Sodalis* endosymbionts display also a large range of genome sizes, from the 0.35 Mb of *Candidatus* Mikella endobia (Husnik and McCutcheon, 2016) to the *circa* 4.5 Mb of *S. glossinidius* or *S. pierantonius*, and with several genome sizes in the range of 1-2 Mb (Santos-Garcia et al., 2017). This range has also been observed in *Serratia symbiotica*, with genome sizes from 0.65 to 3.86 Mb (Manzano-Marín et al., 2016).

Previous findings by Nováková et al. (2009) placed *Arsenophonus* from *A. dispersus* and *A. duguesii* (a close relative), together with an *Arsenophonus* from *B. tabaci*, in a clade with long branches, which is apart from other *Arsenophonus* of whiteflies. Our phylogenomics confirms these results, and have revealed that ARAD belongs to a different clade from the other two whiteflies’ *Arsenophonus*, ARAF and ARTV. Two scenarios can explain the fact that ARAD and ARAF belong to different clades. In the first, the ancestor of ARAD infected the lineage of *A. dispersus* after its divergence from *A. floccissimus* and later started its shift to a primary symbiosis and its associated genome reduction process. In this case, the process could have taken place for as long as 18 My, the estimated divergence time between these two whitefly species (Santos-Garcia et al., 2015). An alternative scenario is that this association began prior to the divergence of the two *Aleurodicus*, but later, ARAD-type symbiont was replaced in the *A. floccissimus* lineage by another *Arsenophonus* lineage (ARAF). Although both scenarios are possible, the large number of pseudogenes in the ARAD genome suggests that the initiation of the process was relatively recent, as suggested by the first scenario. Still, the second scenario cannot be excluded as symbiont replacement has been documented in many insect lineages (Sudakaran et al., 2017).

As far as we know, *Arsenophonus* ARAD is among the smallest genomes in its genus, between *R. pediculicola* and *A. lipopteni* (Šochová et al., 2017). ARAD is still under a genome reduction process that could end up below 0.4 Mb if the current 54 % coding density is considered. ARAD gene repertoire is clearly a subset of other whiteflies’ *Arsenophonus*. It is totally dependent on the host’ environment, and putatively supplies/complements its host with cofactors/vitamins not produced by *Portiera* and the host. In conclusion, ARAD can be considered a co-primary endosymbiont. On the other hand, ARAF and ARTV are still at the beginning of the genome reductive process (e.g. high number of pseudogenes, inactivation of mobile genetic elements, and virulence/secretion factors). They are heritable facultative endosymbionts with the potential to establish a co-primary symbiotic relationship with *Portiera* and their host. However, as frequently observed, all *Arsenophonus*, independently of their symbiotic status, can be replaced by other bacterium of the same, or different genera, as long as the new comers have similar biosynthetic capabilities (Thao and Baumann, 2004; Russell et al., 2017; Sudakaran et al., 2017).

We hypothesize that the ability to synthesize riboflavin and other vitamins is probably the main reason for the presence of *Arsenophonus* endosymbionts in whiteflies and the evolution of ARAD towards a co-primary symbiont in *A. dispersus*. As indicated above, nutritionally poor diets such as blood and phloem sap, cannot provide insects with all their dietary requirements, including several vitamins. For example, the essentiality of dietary riboflavin was demonstrated in aposymbiotic aphids but not in aphids harboring *Buchnera aphidicola*, which is able to produce riboflavin (Nakabachi and Ishikawa, 1999). Moreover, *B. aphidicola* of aphids from the subfamily Lachninae already lost the genes encoding the pathway in the Lachninae ancestor. This led to the establishment in this group of a new association with an additional bacterium, capable of producing riboflavin. In most lineages, this is the co-primary *S. symbiotica*, a facultative endosymbiont in other aphid lineages (Manzano-Marín and Latorre, 2014). The analyses of the co-symbiosis in species of Lachninae revealed a complex system with *S. symbiotica* endosymbionts harboring a very small genome (in *Tuberolachnus salignus*), intermediate genome sizes (in some species of the genus *Cinara*), or replaced by other endosymbionts belonging to different genera, such as, *Sodalis*, *Erwinia*, or *Hamiltonella*, among others (Manzano-Marín et al., 2016, 2017; Meseguer et al., 2017). The essential role riboflavin biosynthesis might play in establishing symbiotic relationships has also been demonstrated in several blood-feeding arthropods harboring different endosymbionts, such as *A. lipopteni* and its host the biting fly *Lipoptena cervi* (Nováková et al., 2016), *Wolbachia* associated with the bedbug *Cimex lectularius* (Moriyama et al., 2015), and *Coxiella*-like bacteria found in several ticks (Gottlieb et al., 2015).

Although, thiamine, pantothenate, coenzyme A, niacin, pyridoxal and biotin can be found in the phloem (free or bounded to transporter proteins) (Pirson and Zimmermann, 1975), their concentrations might be not sufficient for many phloem-feeding species. This could explain why the capability to produce some of these compounds has been kept in bacterial endosymbionts while it has been lost in others. However, the production of one or several B vitamins (thiamine, riboflavin, pyridoxal, folate, and biotin) and *de novo* lipoate by whiteflies’ *Arsenophonus* seems to be the main contribution of these endosymbionts to their hosts, allowing them to feed on the limited diet present in the phloem (Dale et al., 2006; Nováková et al., 2015, 2016; Mao et al., 2017). Regarding biotin biosynthesis, the bacterial origin of *bioAB* and *bioD* was reported by Luan et al. (2015) and Ankrah et al. (2017). We show here that not all whiteflies seem to have the horizontally transferred *bioAD* genes, although the rest of the *bio* operon homologs, including the *bioB*, are contained. This suggest that whiteflies acquired, by several horizontal gene transfer events, the ability to produce biotin. This is expected to produce a gradual loss of the biotin pathway in secondary endosymbionts of the different whiteflies lineages, at least from the Aleyrodinae subfamily, as this group harbors the full *bio* operon. Indeed, *Cardinium hertigii* infecting *B. tabaci* has recently lost its ability to produce biotin, raising the possibility that biotin is provided by the co-present *Hamiltonella* endosymbiont or by the host metabolic potential (Santos-Garcia et al., 2014b; Rao et al., 2015). As an alternative, this can also lead to intricate complementation patterns as shown in the *Arsenophonus* from *Aleurodicus*. It is important to notice that supplementation of vitamins was the role proposed for *Hamiltonella* in *B. tabaci* (Rao et al., 2015) and the results reported here support a similar function for *Arsenophonus*. Interestingly, these two endosymbionts are never found together in the same individual (Gottlieb et al., 2008; Skaljac et al., 2010, 2013; Marubayashi et al., 2014). It could be possible that the two endosymbionts might compete for the same host resources to produce the same compounds, undermining the unique benefit to the host if harbored together due to the cost associated of maintaining high endosymbionts loads (Ferrari and Vavre, 2011).

Finally, and in contrast to *Arsenophonus*, *Wolbachia* from *Aleurodicus* species only produces riboflavin and, probably, folate. It has been proven that the bedbug *C. lectularius* requires the riboflavin produced by *Wolbachia* for its proper development. Indeed, the riboflavin pathway is conserved among *Wolbachia* species infecting different insects, suggesting that it may provide a fitness benefit to the infected host (Moriyama et al., 2015). However, it is yet to be determined if *Wolbachia* can be considered as a general mutualistic symbiont (at the metabolic level) in whiteflies, like *Hamiltonella* or *Arsenophonus*. First, while *Hamiltonella* and *Arsenophonus* are restricted to bacteriocytes, *Wolbachia* is found in different tissues (Gottlieb et al., 2008; Skaljac et al., 2010, 2013; Marubayashi et al., 2014). While the first pattern is more characteristic of hemipteran mutualistic endosymbiosis, the second can be found in endosymbionts with a wide range of symbiotic interactions, including parasitism. Second, in most reported cases of whiteflies harboring *Wolbachia*, the individual insects usually harbor in addition *Arsenophonus* or *Hamiltonella* (Skaljac et al., 2010, 2013; Kapantaidaki et al., 2014; Marubayashi et al., 2014; Zchori-Fein et al., 2014). The few cases in which *Wolbachia* was found to be the unique secondary endosymbiont should be handled with caution, as they may simply result from failure of the general PCR primers to amplify other symbionts species/strains (Augustinos et al., 2011).

In conclusion, the loss of the genes encoding the enzymes for the synthesis of vitamins in the ancestral *Portiera*, many million years ago, likely generated the requirement of a co-symbiont in whiteflies. *Arsenophonus* species seem to be the most common co-symbiont. In the lineage of *A. dispersus*, the harbored *Arsenophonus* lineage presents a highly reduced genome content. This *Arsenophonus* lineage is, or is in the process of becoming, a co-primary symbiont which putatively supplies/complements its host with cofactors/vitamins that are not produced by its co-partner *Portiera*.

### Author Contributions

DSG and FJS conceived the study. DSG and FJS performed bioinformatics analysis. KJ performed phylogenetics analysis. All authors analyzed and discussed the data. DSG and FJS drafted the manuscript with inputs from AL, AM, EZF, SF and SM. All authors participated in the revision of the manuscript.

### Funding

This work was supported by the grants PROMETEOII/2014/065 (Conselleria d’Educació, Generalitat Valenciana, Spain) to AM, BFU2015-64322-C2-1-R (co-financed by FEDER funds and Ministerio de Economía y Competitividad, Spain) to AL, and by the ISRAEL SCIENCE FOUNDATION (Israel) grants No. 1039/12 to SM and No. 484/17 to SF and EZF.

#### Acknowledgments

DSG is recipient of a Golda Meir Post Doctoral Fellowship at the Hebrew University of Jerusalem. The authors acknowledge Francisco J. Beitia and Estrella Hernández Suárez for the whiteflies samples.

### Supplemental Data

The Supplementary Material for this article can be found online at: Supplementary Material

### Data Availability Statement

The *Arsenophonus* (ERZ502704-6) and *Wolbachia* (ERZ502707-8) annotated genomes and the whole genome shotgun libraries (ERR2532344-45 and ERR2534068-71) analyzed for this study have been deposited at the European Nucleotide Archive (ENA) under the project number PRJEB26014. The generated Pathway Tools databases and the blastx results of the genomic reads similar to *Bemisia tabaci* key cofactors/vitamins metabolic genes can be found at 10.6084/m9.figshare.6142307.

